# Real-time monitoring of attenuated cytomegalovirus using Raman spectroscopy allows non-destructive characterization during flow

**DOI:** 10.1101/2024.05.08.593031

**Authors:** Shreya Milind Athalye, Murali K. Maruthamuthu, Ehsan Esmaili, Miad Boodaghidizaji, Jessica Raffaele, Vidhya Selvamani, Joseph P. Smith, Tiago Matos, Richard R. Rustandi, Arezoo M. Ardekani, Mohit S. Verma

**Author notes:** Authors contributed equally (Shreya Milind Athalye, Murali K. Maruthamuthu, Ehsan Esmaili).

## Abstract

Real-time monitoring of viral particles can have a crucial impact on vaccine manufacturing and can alleviate public health by supporting continuous supply. Spectroscopic methods such as Raman spectroscopy can provide rapid and non-invasive measurements. Here, we have developed a Raman spectroscopy-based tool to monitor the quality and quantity of viral particles in a continuous flow set-up. The attenuated human cytomegalovirus (CMV) is characterized across a wide range of concentrations (4.50 x 10^9^ to 2.90 x 10^11^ particles/mL) and flow rates (100 µm/s to 1000 µm/s) within a square quartz capillary. This process analytical technology (PAT) tool enables the detection of viral particles even at high flow rates such as 1000 µm/s. Sodium dodecyl sulfate-polyacrylamide gel electrophoresis (SDS-PAGE) and dynamic light scattering (DLS) demonstrated that the samples maintain their integrity even after laser exposure, reiterating the non-invasive nature of Raman spectroscopy. To the best of our knowledge, this is the *first* report on characterizing CMV particles using Raman spectroscopy. We have also demonstrated the limit of detection (LoD) (2.36 x 10^10^ particles/mL) for CMV particles in continuous flow (via the Raman spectroscopy method) by addressing the effect of flow rate, concentration, and integrity of samples. This technology could enhance our understanding of the quality control in bio-manufacturing processes required in vaccine production.

## 1. Introduction

Vaccines remain the most effective method to control both endemic and pandemic infections. Their production, however, still frequently relies on the same type of technologies developed decades ago. New processing technologies offer attractive advantages, including increased efficiency and productivity, and are now slowly being implemented across vaccine production.^1–6^ As evidenced by the severe acute respiratory syndrome coronavirus 2 (SARS-CoV-2) pandemic, the rapid, robust deployment of emergent vaccines at a considerably high dosage requires the transformation of the existing technologies and strategies for vaccine production.^7^ Besides the faster deployment, the need for cost-efficient processes has brought forward new downstream alternative purification approaches.

Among some of these new approaches, novel strategies for process intensification, such as continuous or semi-continuous systems, offer the potential for tremendous process efficiency improvements and increased robustness while simultaneously offering a reduction of footprint and cost of goods.

Continuous manufacturing (CM) has been implemented across several industries, such as oil, chemicals, synthetic fibers, fertilizers, power, natural gas, and automobiles.^8^ CM has recently gained momentum in the pharmaceutical industry, as noted by the first oral drug produced (Orkambi®) via CM being granted Food and Drug Administration (FDA) approval in 2015 and multiple additional FDA-approved CM-based drugs since then.^9,10^ Building on these approaches, interest in employing CM throughout the pharmaceutical and other industrial communities continues to rise, especially given recent guidance drafts from the FDA for facilitating CM implementation with targeted economic models for driving various pharmaceutical pipelines forward via CM.^8,11–13^ A variety of recent developments in process analytical technology (PAT) have allowed process teams to monitor their products in realtime throughout various production steps.^8,11–22^ Although the latest developments are exciting,^23^ CM is still an untapped resource within the large molecule pharmaceutical communities, including for viruses and virus-like particles. A key reason for this gap is a lack of control strategies beyond automation and process design. As such, the need for new PAT within the CM of large molecules is critical.

Most of the current analytical techniques for the quantification and characterization of viral particles—such as quantitative polymerase chain reaction (qPCR), droplet digital polymerase chain reaction (ddPCR), dynamic light scattering (DLS), flow virometry,^14^ and quantitative capillary western blot,^15,16^—are still employed as at-line or offline methods. Therefore, improvement is still needed to ensure precise real-time, *in situ* monitoring of the quality and quantity of the viral particles in continuous manufacturing.^17^

There has been a rapid development in optical techniques and plasmonic biosensing tools to analyze biomedical samples.^24–28^ This development includes a variety of complex techniques such as Raman spectroscopy and surface plasmon resonance to detect and amplify the chemical signature of bio-samples due to weak inelastically scattered light and local enhancement of electric properties of plasmonic structure (e.g., Au or Ag).^25–27^ As a result, there have been numerous studies on the detection of different viruses and cells such as Human immunodeficiency viruses (HIV,) cancer cells, H1N1 Influenza, and the novel coronavirus SARS-CoV-2.^26,27^ These techniques involve the use of nanomaterials and biological probes to facilitate fast quantification of viral particles. Non-uniform adsorption of target virus particles and resulting non-linear signal enhancement due to local high concentration limits its translation into *in situ* continuous monitoring of viral particles. Biological probes used to increase the specificity and sensitivity of viral particle detection usually function as reporters generating optical signals, limiting their use in the qualitative monitoring of vaccine production.

Raman spectroscopy provides detailed molecular fingerprints of viral particles even in an aqueous state, enabling qualitative and quantitative in situ monitoring in manufacturing processes.^18,29^ Previously, we have implemented this technique to detect and categorize bacteria, fungi, and mammalian cells combined with a deep learning algorithm.^30– 32^ Building on our earlier work, we now implement Raman-based viral quantification in a continuous flow and characterize the viral loads at different concentrations and flow rates.

We have developed a Raman spectroscopy-based PAT tool that can facilitate continuous monitoring of attenuated CMV at different concentrations (4.50 x 10^9^ to 2.90 x 10^11^ particles/mL) at various flow rates (100 µm/s to 1000 µm/s) within a square quartz capillary set-up. In this study, we demonstrate the following advancements: i) We characterized CMV particles using Raman spectroscopy and identified peaks for CMV viral particles that are distinct from the background peaks coming from buffer, ii) We demonstrated the detection of the CMV particles in flow conditions that are similar to the industrial operating conditions, iii) We identified limit of detection (LoD) for multiple Raman peaks that can be used for continuous monitoring of the CMV particles, iv) We demonstrated that our technology is non-invasive and maintain the integrity of samples.

## 2. Materials and Methods

### 2.1 Viral production and purification

The vaccine virus was produced in human retinal pigment epithelium cells (ARPE-19)^33^ grown on Cytodex-1 ® microcarrier using a proprietary growth medium in a stirred-tank bioreactor. After sufficient cell growth was achieved, the bioreactor was 80% medium exchanged, followed by viral inoculation. At 14 days post-infection, the cell culture was harvested by removing the microcarrier beads through a strainer bag and subsequently clarified through Sartobind GF+ 1.2 µm filter. We further used this clarified-viral harvest for subsequent chromatographic studies.

During purification, single membrane runs for scouting experiments were performed with an Äkta Pure system (Cytiva, Upsala) using 10mM histidine (addition of 150 mM sodium chloride (NaCl), pH 7) equilibration buffer at 10 mL/min on 3 mL Sartobind Q nanomembranes, for a total of 5 column volumes (CV). A wash step was performed for 15 CV at 180 mM NaCl and 80 mM histidine, with a pH of 6. A 15 CV gradient step to 1 M NaCl (in 25 mM Histidine buffer, pH 7) was applied for elution. In addition, alternative loads were performed using a wash buffer as a mix-loading buffer. All collected fractions were analyzed off-line by Sodium dodecyl sulfate-polyacrylamide gel electrophoresis (SDS-PAGE) and Apogee to estimate yields. For concentration, tangential flow filtration was performed using a 750 kDa hollow fiber (Cytiva, Upsala).^34^

### 2.2 Fabrication of capillary tube set-up for viral samples under flow

A single capillary device was fabricated to characterize the ability of the Raman microscope to detect viral particles in a continuous flow set-up. We used a customized crystal-clear fused quartz square capillary tube (Friedrich & Dimmock, Inc, NJ, USA) with a size of 2 mm× 2 mm × 50 mm (0.75 mm wall thickness). The square capillary tube was bonded on a glass slide using UV epoxy resin (60-7105, Epoxies Inc, RI, USA). Both ends of the quartz capillary were connected to plastic tubes (ID 2mm, 5233k112, McMaster-Carr, IL, USA), and the ends were sealed with UV epoxy resin. The viral samples were pumped into the capillary using a syringe pump (70-3007, Harvard Apparatus, MA, USA).

### 2.3 Raman analysis on a square capillary tube in static and flow conditions for viral samples

To initially characterize the performance of our Raman microscope with liquid samples, we tested liquid attenuated CMV and obtained Raman spectra at different conditions (laser power and acquisition time). The capillary tube setup mentioned above was used for the static and flow condition study. The capillary tube set-up, along with the syringe pump, mimicked an on-line flow set-up. The attenuated CMV was initially tested at static conditions (0 µm/s) and with a flow rate from 100 µm/s to 1000 µm/s (24-240 µL/min). The above flow rate range is based on the viral vaccine production platform (320 cm/h = ∼900 µm/s). A syringe pump provided the different flow rates inside the capillary tube. The attenuated CMV was purged in the capillary set-up, and the viral sample was analyzed immediately using the Renishaw inVia^TM^ Qontor® confocal Raman microscope (Renishaw plc, Wotton-under-Edge, UK). We used a 785-nm excitation laser with 50% (∼150 mW) power, objective 20X, and 10 s acquisition time per measurement. Objective 20X has a numerical Aperture (NA) of 0.40, a working distance of 1.15 mm, a depth of focus of 9.81 µm, and a laser spot of 2.4 µm diameter with pinhole out. The collected Raman spectra ranged from 100-3200 cm^-1^ with a spectral resolution of 1 cm^-1^. We collected 20 technical replicates using the map acquisition step, which allowed us to collect the Raman spectrum from different points/XY-coordinates of the sample. To accurately distinguish the Raman peaks from attenuated CMV, we prepared a synthetic buffer with all excipients except the viral particles and used it as a background control. We used internal Si measurement as an internal standard to calibrate. Before every experiment, we calibrated the Raman microscope using a silicon (Si) sample as an internal standard to assess any alignment drift in the system. Here, we collected Raman spectrum for Si and fitted the spectrum (peak 520 cm^-1^). We confirmed if the peak position and intensity matched with the previous spectrum for the same sample at the same conditions and adjusted offset if the peak position was observed beyond ±0.5 cm^-1^ (Section A, Table S1). To study the sensitivity of our Raman microscope with the liquid viral samples, we tested different dilutions of viral samples (attenuated CMV in the concentration range of 4.50 x 10^9^ to 2.90 x 10^11^ particles/mL) in the same setup. The raw data for Raman spectra were obtained using WiRE™ 5.5 software.

### 2.4 Raman Data processing

Raman spectral data processing pipeline involved baseline correction algorithm, normalization, and area under the curve calculation. The raw data were baseline corrected using the adaptive iteratively reweighted Penalized Least Squares (airPLS) algorithm^35^ and Python script adapted by Renato Lombardo^36^ of the University of Palermo. The Python script was modified to remove cosmic rays and normalize Raman spectrum using Linear Normalization. The Python script also facilitated the area under the curve calculations.^37^ A two-sample t-test was used to identify distinct peaks from the buffer. A linear fit was applied to the area under the curve calculations to obtain LoD plots. The Limit of Blank (LoB) was calculated using the area under the curve for buffer (viral particle concentration: 0 particles/mL). The formula for the Limit of Blank calculations is as follows:

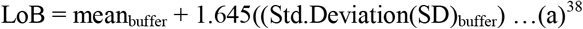

The LoD for each Raman peak was calculated as follows:

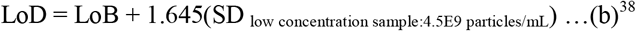

All plots were plotted using OriginPro 2022 (OriginLab Corporation, Northampton, MA, US) software.

### 2.5 Dynamic light scattering analysis to check the effect of laser exposure

The effect of Raman laser exposure (785-nm excitation laser with 50% (∼150 mW) power with 10 s total acquisition time per scan) on the viral particles was characterized using the dynamic light scattering (DLS) technique. This analysis will evaluate the effect of laser exposure on the size of the viral particles. The CMV samples in the concentration range of 4.50 x 10^9^ to 2.90 x 10^11^ particles/mL were used to evaluate this effect. Before and after the laser exposure, the size of the viral particles was analyzed using a disposable plastic micro cuvette (ZEN0040, Malvern, MA, USA) in the Nano-S Zetasizer (Malvern, MA, USA).

### 2.6 SDS-PAGE analysis before and after laser exposure

The functional impact of Raman laser exposure on the viral samples was evaluated. SDS-PAGE was performed to analyze any changes during the laser exposure to the protein samples during Raman spectra acquisition. SDS-PAGE was performed using 4–15% Mini-PROTEAN® TGX™ Precast Protein Gel (#4561084, Bio-Rad Laboratories, Inc, USA). Before and after the laser exposure, the samples were collected, reduced in sample buffer containing 100 mM Dithiothreitol (DTT), and heated at 70 °C for 10 min. The two CMV (before and after laser exposure) samples were loaded into gels with equal volume (20 µL). Electrophoresis was performed with constant voltage at 125 V for one hour. The gel was fixed with 12% trichloroacetic acid, washed with water, stained for >2 hours with Bio-Safe™ Coomassie Stain (#1610786, Bio-Rad Laboratories, Inc, USA), and de-stained in water for > 2 hours. The gel was scanned by UVP ChemStudio touch (model 695, analytikjena, USA).

## 3. Results and Discussion

### 3.1 Raman spectroscopy-based non-invasive tool for real-time monitoring

An effective process monitoring approach involves monitoring the state of the process, detecting transient disturbances, and enabling continuous operation through critical control of material collection/diversion and real-time release testing. One crucial aspect that could facilitate process monitoring is Sampling frequency. Sampling frequency should be fast enough to detect rapid changes and frequent enough to process drift that could facilitate the harvest/transfer of the product to the next step. It should also enable trend analysis and statistically sound data to make informed quality controls.^39^ Here, we followed a methodology that enabled real-time monitoring of CMV particles through fast and frequent sampling under different operating conditions, such as static vs. flow conditions. We quantify the concentration of CMV samples under static and flow conditions. Figure 1 overlays the study schematic involving sample preparation, analytical device (substrate for Raman), and confocal Raman microscope. To determine the LoD of this system, we prepared different concentrations of CMV particles by serial dilution (Figure 1 a). Figure 1b indicates the analytical device comprising a capillary set-up, which stimulated the required flow conditions for analysis. All experiments were performed on the confocal Raman microscope, as shown in Figure 1c.

**Figure 1.**
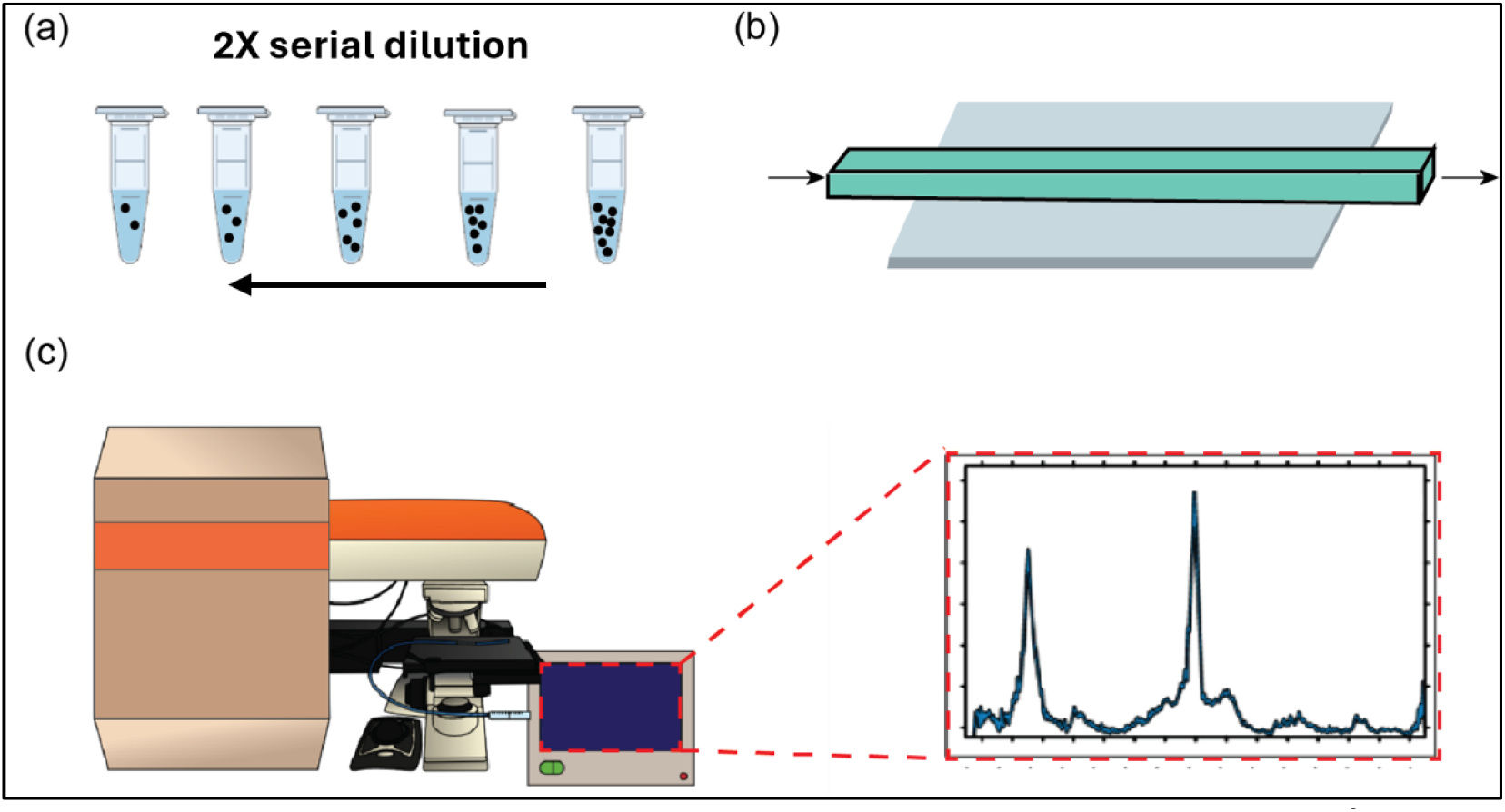
Schematic of the set-up. (a) Preparation of CMV viral dilutions from 4.50 x 10^9^ particles/mL to 2.90 x 10^11^ particles/mL. (b) Square quartz capillary tube with inlet and outlet for flow. (c) Raman imaging with the viral flow using inVia™ Quontor® confocal Raman microscope. Inset shows on-line Raman data collection.

### 3.2 Raman spectroscopy for CMV particles

CMV is an enveloped herpes virus, with multiple glycoproteins embedded on an outer lipid bilayer envelope and a double-stranded linear DNA.^40,41^ The surface glycoproteins are essential to entering host cells and, thus, are important targets for neutralizing antibodies.^42^ Current CMV vaccine strategies focus on virus-derived, recombinant subunit proteins or viral vectors. Identifying and characterizing these virus-like particles that can accurately predict various quality attributes, including concentration and purity, is valuable in translation to continuous manufacturing. We employed Raman spectroscopy to characterize CMV particles in capillary set-up in static conditions. We selected Raman peaks for CMV particles that were distinct from the background peaks from the buffer and capillary setup. The Raman spectrum (Figure 2) of CMV particles and background are displayed as normalized Raman Intensity vs Raman shift.

**Figure 2.**
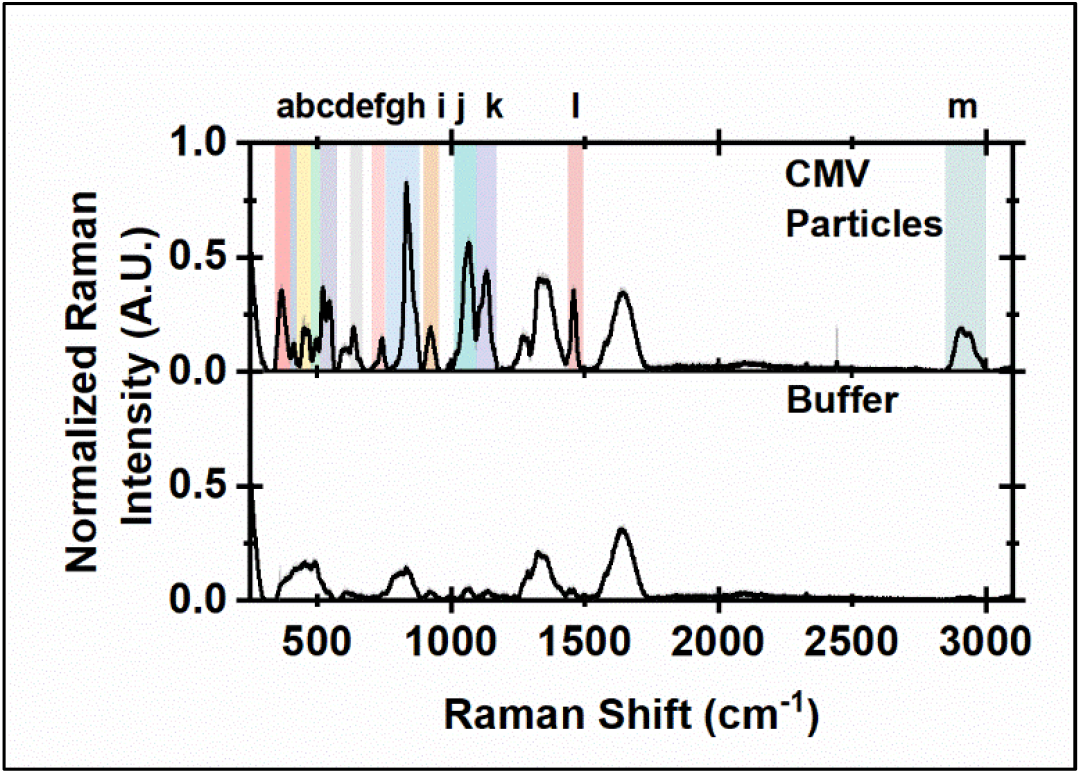
Normalized Raman spectra for attenuated cytomegalovirus (CMV) viral particles in static condition (concentration: 2.90 x 10^11^particles/mL) (Range for Raman peaks is as follows-a: 343-402 cm^-1^, b: 402-425 cm^-1^, c: 425-476 cm^-1^, d: 476-510 cm^-1^, e: 510-575 cm^-1^, f: 627-669 cm^-1^, g:704-756 cm^-1^, h: 760-884 cm^-1^, i: 899-954 cm^-1^, j: 1011-1095 cm^-1^, k: 1095-1174 cm^-1^, l: 1438-1497 cm^-1^, m: 2850-3000 cm^-1^)

We identified 13 peaks (labeled a-m in Table 1) that are distinct from the background spectra and correlate strongly to the CMV particles (Figure 2, colored bars). Table 1 describes the functional group assignment and interpretations of the composition for each peak. The peak centered at 2900 cm^-1^ corresponds to the C-H stretch, which is important to indicate the capsid structure of the virus.^29^ Other peaks centered at 921, 1072, and 1127 cm^-1^ correspond to C-C-N and C-N stretches, which correspond to the nucleic acids and some amino acids in the viral particle. C-S stretch region was observed at 639 and 744 cm^-1^, which could be assigned to aromatic amino acids such as Phenylalanine and Tyrosine. Previously, characterization of different viruses, including HIV, Ecovirus 1, Bacteriophage PRD1, and SARS-CoV-2, also observed most of the above peaks.^43–47^

**Table 1:** Raman peak frequencies, range of the peak, tentative functional group assignments for the Raman peaks, and Interpretations for CMV characterization.

| Sr. no. | Raman peak Frequency ($\text{cm}^{-1}$) | Range of the peak ($\text{cm}^{-1}$) | Functional Group Assignment<sup>29</sup> | Interpretations<sup>29,49</sup> |
| --- | --- | --- | --- | --- |
| **a** | 369 | 343-402 |  |  |
| **b** | 416 | 402-425 |  |  |
| **c** | 461 | 425-476 |  |  |
| **d** | 496 | 476-510 | S-S stretch |  |
| **e** | 525, 547 | 510-575 | S-S stretch |  |
| **f** | 639 | 627-669 | C-S stretch | Phenylalanine, Tyrosine |
| **g** | 744 | 704-756 | C-S stretch | Thymine, Tyrosine, Adenine |
| **h** | 836 | 760-884 | (O-P-O) symm. Stretch | Sugar-phosphate symmetric stretch |
| **i** | 921 | 899-954 | (C-C-N) symm deformation | α-helical skeletal |
| **j** | 1072 | 1011-1095 | C-N stretch | Phenylalanine, Nucleic acids |
| **k** | 1127 | 1095-1174 | C-N stretch | Phenylalanine, Tyrosine |
| **l** | 1463 | 1438-1497 | Aromatic ring /CH<sub>2</sub> scissoring/ CH<sub>2</sub>/CH<sub>3</sub> deformation | Adenine |
| **m** | 2900 | 2850-3000 | CH stretch |  |

**Table 2:** Two sample t-tests for different dilutions of attenuated cytomegalovirus (CMV) viral particles in static conditions indicate that the Raman spectral for peak 2900 cm^-1^ of the viral particles is significantly different from the background spectra.

| Raman Shift ($\text{cm}^{-1}$) | Concentration (Particles/mL) | Normalized Raman Intensity (A.U.) | | S/N ratio |
| --- | --- | --- | --- | --- |
|  |  | Avg (n=20) | SD(n=20) |  |
| **2907.18** | $2.90 \times 10^{11}$ | 0.192 | 0.009 | 19.42 |
| | $1.50 \times 10^{11}$ | 0.098 | 0.007 | 13.99 |
| | $7.30 \times 10^{10}$ | 0.054 | 0.007 | 7.76 |
| | $3.60 \times 10^{10}$ | 0.036 | 0.007 | 4.89 |
| | $1.80 \times 10^{10}$ | 0.024 | 0.007 | 3.37 |
| | $9.10 \times 10^9$ | 0.016 | 0.004 | 4.00 |
| | $4.50 \times 10^9$ | 0.018 | 0.008 | 2.15 |

**Table 3:** Signal-to-noise ratio for different dilutions of attenuated cytomegalovirus (CMV) viral particles in static conditions (peak: 2900 cm^-1^).

| Concentration (Particles/mL) | The area under the curve for peak $2900\text{ cm}^{-1}$ | | |
| --- | --- | --- | --- |
| | Avg(n=20) | SD(n=20) | p-value ($\alpha=0.05$) |
| **$2.90 \times 10^{11}$** | 15.59 | 0.79 | 9.71E-44 |
| **$1.50 \times 10^{11}$** | 7.82 | 0.46 | 7.74E-39 |
| **$7.30 \times 10^{10}$** | 4.55 | 0.15 | 6.02E-42 |
| **$3.60 \times 10^{10}$** | 3.22 | 0.28 | 1.34E-24 |
| **$1.80 \times 10^{10}$** | 2.73 | 0.19 | 4.68E-24 |
| **$9.10 \times 10^9$** | 1.82 | 0.12 | 1.16E-07 |
| **$4.50 \times 10^9$** | 2.04 | 0.18 | 8.78E-12 |

The Raman peak at 2900 cm^-1^ in the C-H stretching region was the most prominent. The C-H stretching region is attributed to the diversity of alkyl groups but also indicates the stability of amino acids. Any changes in the structure or composition of the amino acids could affect the Raman peak in the C-H stretching region.^48^ Thus, the C-H region can help us interpret the concentration and stability of the viral particles. We selected this peak for further studies.

### 3.3 Characteristics of C-H stretching region of CMV

We further characterize the Raman peak at 2900 cm^-1^ to determine its suitability in the quantification of the CMV viral particles. The Raman peak at 2900 cm^-1^ showed a uniform trend of decreasing Raman intensity with decreasing concentrations under static conditions (Figure 3a). We used Gaussian fit to deconvolute the peak of the interest (2900 cm^-1^) (Figure 3b). The Gaussian fit identified three Raman peaks centered at A) 2903.23 cm^-1^, B) 2942.92 cm^-1^, C) 2976.93 cm^-1^. Raman peaks centered at 2903.23 cm^-1^ mainly come from CH_2_ modes, and 2942.92 cm^-1^ and 2976.93 cm^-1^ primarily come from the CH_3_ modes.

**Figure 3.**
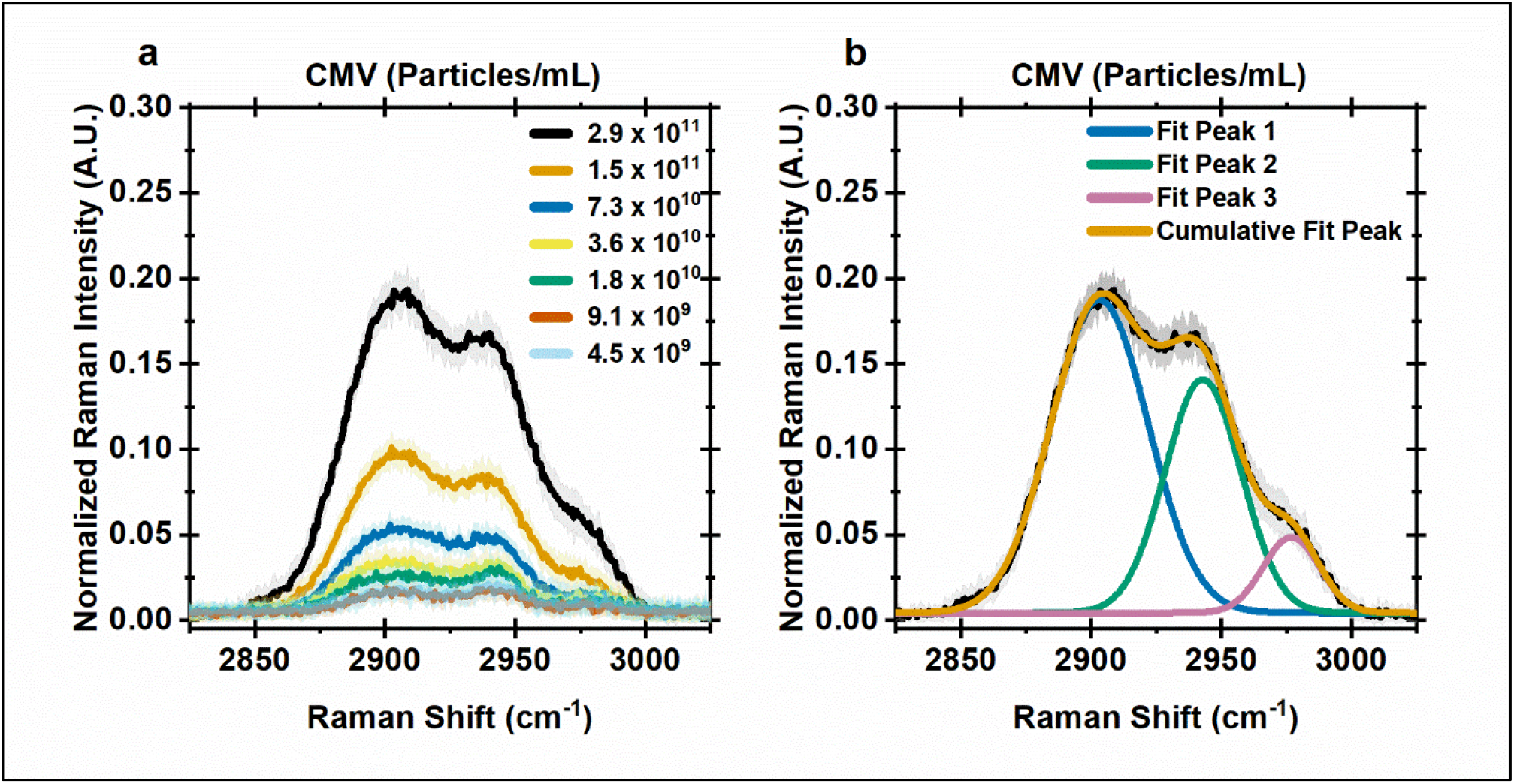
Normalized Raman spectra for attenuated cytomegalovirus (CMV) viral particles in static conditions. (a) Raman spectra for 2900 cm^-1^ peak all the dilutions in the static condition. (b) Deconvolution with Gaussian functions of the Raman spectra of CMV (concentration:2.90 x 10^11^ particles/mL) for the three peaks centered at A) 2903.23 cm^-1^, B) 2942.92 cm^-1^, C) 2976.93 cm^-1^. Before applying the deconvolution process, the fluorescence background has been subtracted.

Biologically, many amino acids have shown more than one peak in the C-H stretching region (2800-3100 cm^-1^). As C-H stretching vibrations are affected by the chemistry of the functional group and its surrounding environment, overlapping of the vibrations can occur, resulting in the broadness of peaks. Thus, the composition and concentration of amino acids in the viral sample could affect the C-H stretching region. Gaussian peaks in the region of 2901-2909 cm^-1^ are observed in Arg (2905 medium-strong), Ala (2906 weak), Ser (2906 medium-strong), Ile (2911 strong), Val (2913 strong), Leu (2913 strong). The region, 2934-2945 cm^-1^, could be from Arg (2939 s), cystine (2941 s), Leu (2941 s), Lys (2941 s), Ile (2948 s), Phe (2951 m), Tyr (2951 m), Glu (2952 s), Thr (2952 s), Val (2953 s), Pro (2954 s), His (2955 s), Asp (2955 s), Asn (2956 s). The Region, 2956−2977 cm^-1^ could involve Ala (2961 s), cystine (2963 m), Cys (2966 s), His (2969 s), Ser (2970 s), Leu (2976 s), Trp (2976 s) Gly (2981 s), Lys (2981 m), Ile (2984 m-s), Val (2985 s).^25,48^ This reiterates the importance of monitoring the C-H region to detect the stability of the viral particles. In the following steps, we moved to detect this Raman peak (2900 cm^-1^) in flow conditions.

### 3.4 Raman spectroscopy for CMV particles in flow conditions

We studied the effect of flow rate from 100 µm/s to 1000 µm/s on the Raman spectra for different concentrations (4.50 x 10^9^ to 2.90 x 10^11^ particles/mL) of CMV particles. The range of flow rates was based on the viral vaccine production platform (320 cm/h = ∼900 µm/s). To characterize CMV particles in flow, the samples were pumped through the capillary tube setup with different flow rates using the syringe pump. We averaged all the technical replicates of flow rates and obtained linear fit (Figure 4a), which was further used to calculate the Limit of detection (LoD) (see Experimental methods for details).^38^ Figure 4a was plotted on a logarithmic scale (log2) for visualization purposes.

**Figure 4.**
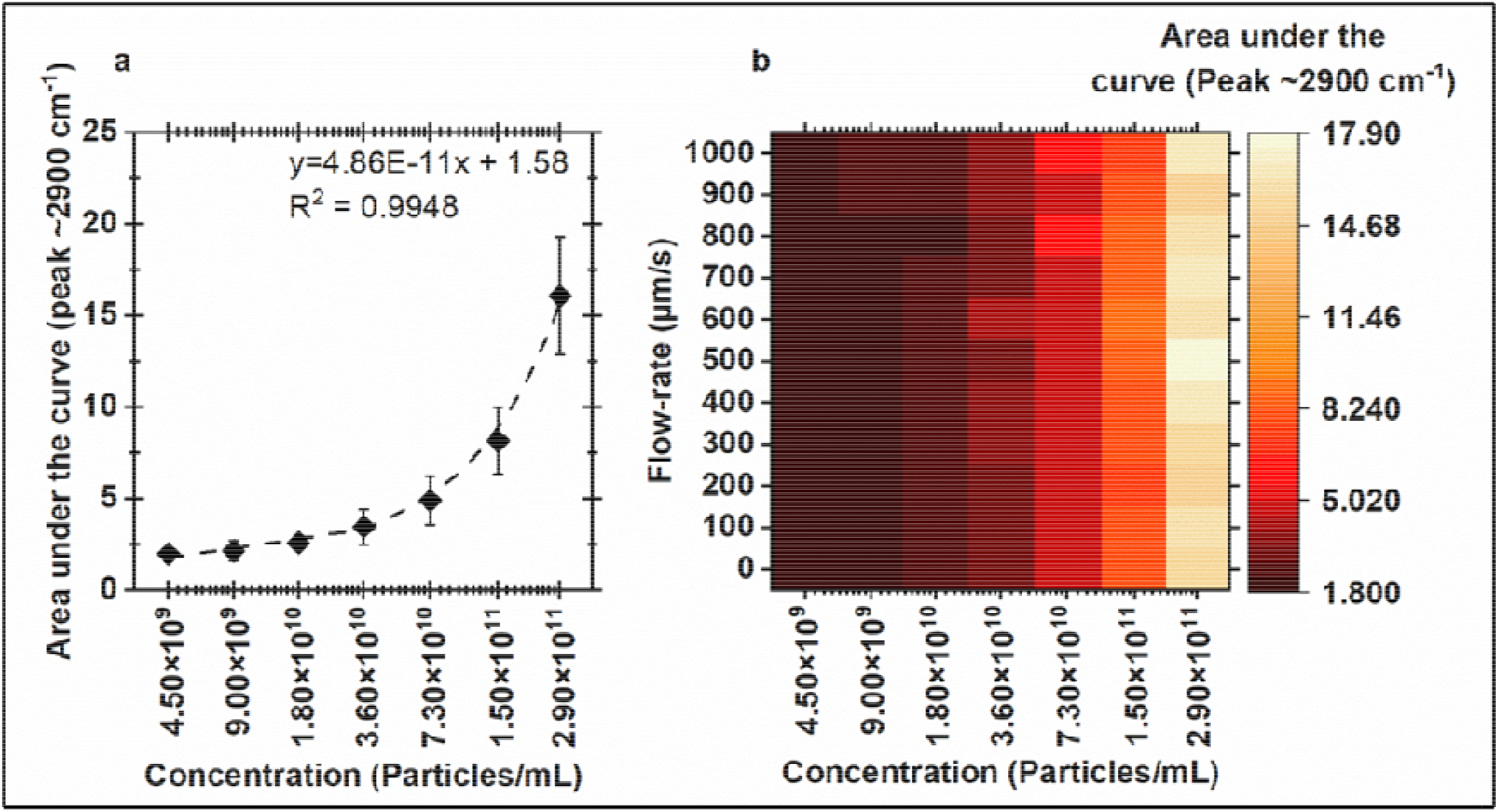
Area under the curve of Raman spectra for 2900 cm^-1^ peak of different dilutions for attenuated cytomegalovirus (CMV) viral particles in flow conditions in the fluid flow setup (Flow rate: 0-1000 µm/s). a) The linear fit for CMV particles: Each point indicates the average area under the curve (n=220), and the error bars indicate the standard deviation (n=220) for the range 2850-3000 cm^-1^. b) Heat map showing the effect of fluid flow (Flow rate: 0-1000 µm/s). Each box indicates the average area under the curve (n=20). Sub-plots a) and b) are represented in a logarithmic scale (log2) for visualization purposes.

The collected Raman spectra were not screened for outliers, and all the technical replicates were used for the analysis. We assumed that viral samples were homogeneous and maintained uniform flow through the capillary. Any non-uniformity in the collection of Raman spectrum was averaged during the 10s of laser exposure time. The calculated LoD for the Raman peak centered at ∼2900 cm^-1^ is 2.36 x 10^10^ particles/mL. LoD calculations for other Raman peaks are shown in Section B, Table S2.

Figure 4b is a heatmap indicating the effect of flow rate for different concentrations as a measure of the area under the curve of Raman peak 2900 cm^-1^. We observed no trend across the flow rates, indicating no significant effect on spectral data. We observed higher deviations in the higher concentrations of the CMV particles (>7.30 x 1010 particles/mL) compared to the lower concentrations (4.50 x 10^9^ to 3.60 x 10^10^ particles/mL). Our results demonstrate that Raman spectroscopy can quantify the CMV particles even at higher flow rates, such as 1000 µm/s. To our knowledge, this is the first demonstration of viral particle detection in flow conditions using Raman spectroscopy. Previously, different PAT tools based on chromatography, DLS, and flow cytometry have been explored in continuous production and monitoring environments for CMV particles.^14,23^ Goldrick et al. used high-throughput Raman spectroscopy in upstream and downstream bioprocessing in the form of at-line and offline analytical tools to eliminate the use of multiple analytical devices. Implementation of the partial least square (PLS) model and Multivariate data analysis (MVDA) facilitated the prediction of multiple process parameters from Raman spectra.^21^ This study could be used as a proof-of-principle and further explored to monitor other process parameters with the help of machine learning and chemometric tools.

### 3.5 Raman spectroscopy is non-destructive

An important factor in developing a PAT tool is to ensure the tool is not destructive to the product (Viral particles /vaccine). Here, the viral particles were exposed to a laser with an excitation wavelength of 785 nm at 50% (∼150 mW) power. We analyzed the viral particles before and after the laser exposure using SDS-PAGE and Western blot to ensure that the laser did not damage the CMV samples. We also determined the effect of laser exposure on particle size using DLS. Based on the DLS result, we found no significant changes in the size of viral particles before and after laser exposure (Figure 5). Preliminary analysis (Section C, Figure S2) has shown SDS-PAGE and western blot for before and after laser exposure (Laser power: 0.5% (∼1.5mW)) does not show any degradation for the critical glycoprotein complexes (gH-gL-UL128 and gH-gL-gO).^16^ Figure 6 indicates that even at higher laser power (50%: ∼150mW), viral particles are stable, and it can be confirmed by comparing SDS-PAGE protein bands in Figure 6. This significant result increases the possibility of adapting this technology as an analytical tool for continuous monitoring of viral particles for vaccine production. We also used COMSOL Multiphysics simulation to calculate the flow velocity and shear stress within the capillary tube (section D, Fig. S3). Based on the calculated Stokes number (ratio of a particle response time to a characteristic time of the flow in the capillary, Stk∼10^-9^), the viral particles follow the fluid streamlines, assuring the uniform distribution of viral particles inside the tube. Also, the range of shear stress inside the capillary is 10^-4^ -10^-3^ Pa, which is lower than typical shear flow experiments involving human CMV samples.^50^ This shows no potential damage to the samples in this range of flow and shear rates.

**Figure 5.**
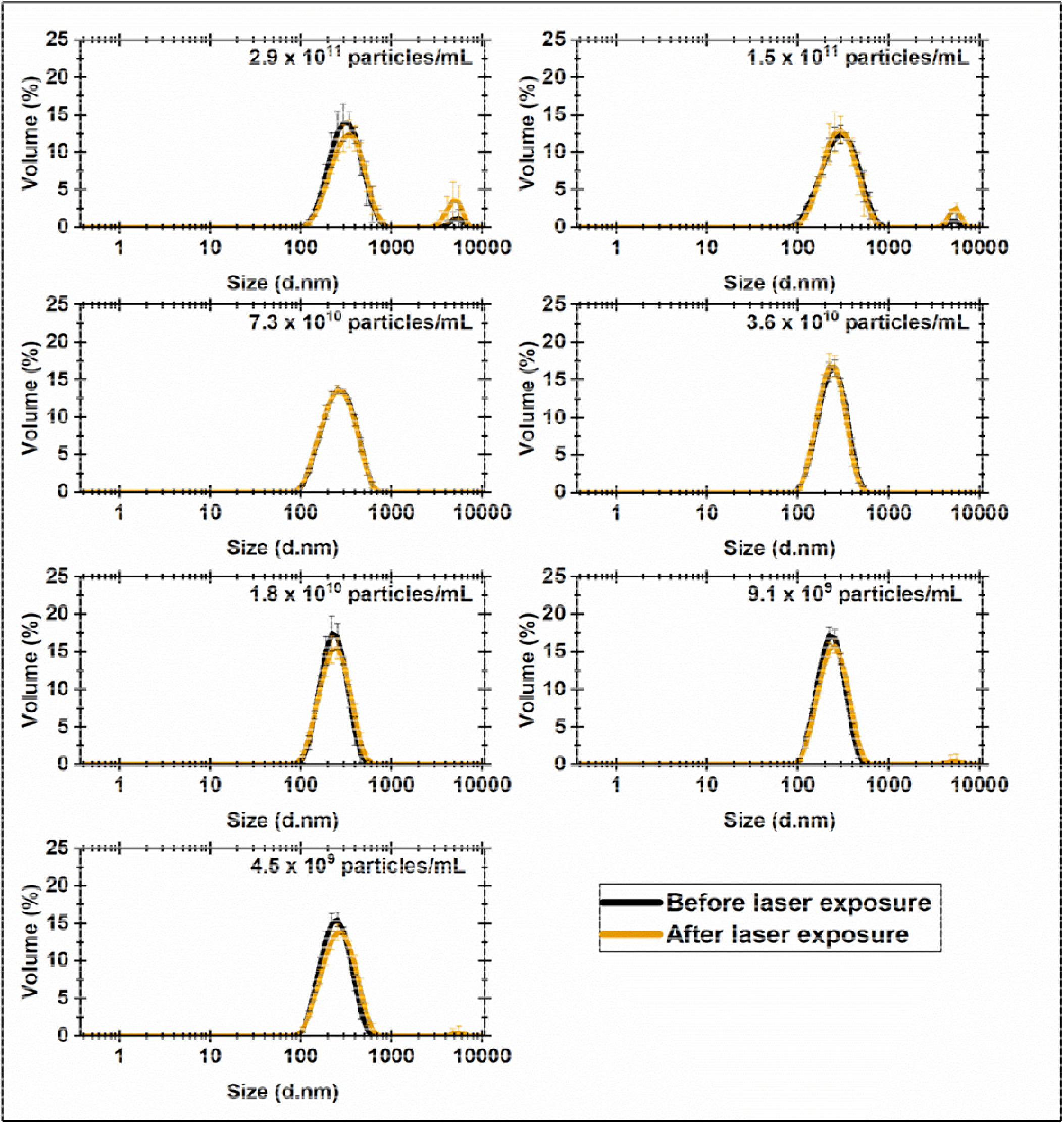
CMV viral particle distribution for different concentrations through DLS analysis. Measurements were done before and after laser exposure (n=5, technical replicates. Each technical replicate is an average of 12 scans).

**Figure 6.**
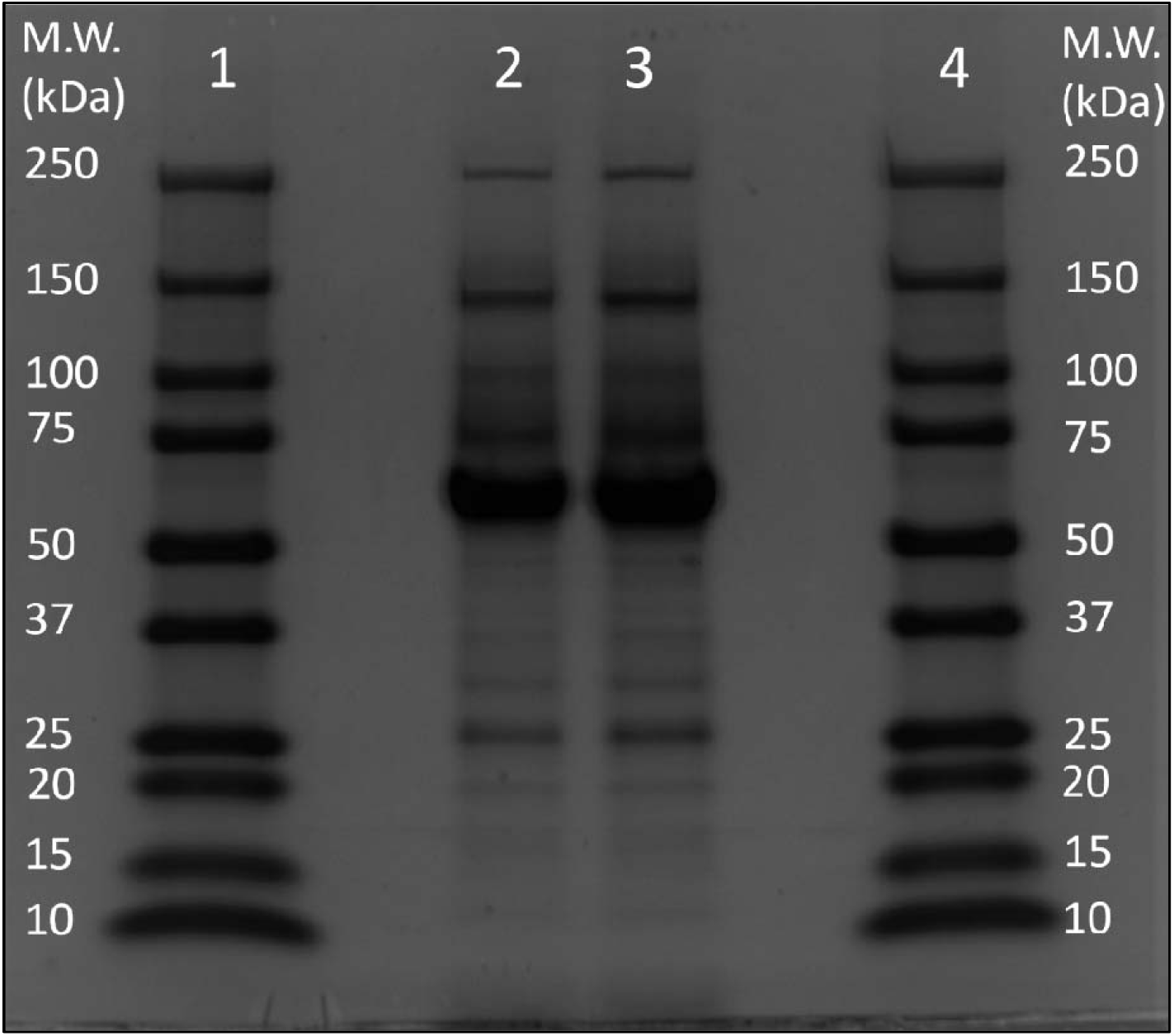
Functional analysis (SDS-PAGE) of the CMV viral particles before and after laser exposure (wavelength: 785nm, Laser power: 50% (∼150 mW), exposure time: 10s, n=20). Lane 2 – before laser exposure and Lane 3 - after laser exposure. Lane 1 and 4 have Precision Plus Protein™ Dual Color Standards, Range: 250-10 kDa, Bio-Rad, #1610374.

## 4. Conclusion

We developed a Raman spectroscopy-based PAT tool that can be applied to on-line analysis of CMV viral particles. Our tool has the following advantages: i) It can be used for qualitative as well as quantitative analysis, ii) it demonstrates its application in static as well as flow conditions similar to industrial vaccine production, iii) It facilitates rapid analysis (∼10s) and thus, can be used for real-time monitoring, iv) It is non-destructive and maintains the integrity of the sample, v) It can easily be applied to other viral particles and bio-processes.

A couple of limitations of the current study are: i) The current study was completed using a non-portable, confocal Raman microscope with high sensitivity; it would be essential to evaluate the performance of this tool with portable Raman probes; ii) Current study evaluates the samples from the downstream stages of the processes which are relatively homogeneous; integration of chemometric tools, multivariate analysis, and machine learning approaches can be explored to improve the sensitivity of this tool for heterogeneous samples.

In future studies, we aim to use microfluidics and acoustic concentration to improve the sensitivity of this Raman spectroscopy-based PAT tool. To our knowledge, this is the first report of a Raman spectroscopy-based PAT tool for monitoring CMV particles. Our current work serves as a proof-of-principle to accelerate the efforts toward PAT development in large molecules.

## 5. Data availability statement

The data sets generated and/or analyzed during the current study are available in the Mendeley data repository at DOI: 10.17632/cs888gdmp4.1

## 6. Acknowledgment

## 7. Author contribution

**Shreya Milind Athalye** contributed the following: design of the experiments (lead), collected and analyzed all the Raman data (lead), writing – original draft (support), writing – formatting (lead), writing – review & editing (lead). **Murali Kannan Maruthamuthu** contributed the following: design of the experiments (support), collected and analyzed all the Raman data (support), writing – original draft (lead), writing – formatting (support), writing – review & editing (support). **Ehsan Esmaili** contributed the following: fabrication of capillary device and fluid analysis (lead) and supported in collecting data and writing the manuscript (support). **Miad Boodaghidizaji** contributed the following: wrote a Python script for data analysis (lead). **Jessica Raffaele** contributed the following: conducting characterization experiments for viral particles before and after laser exposure (support) and review the manuscript (support). **Vidhya Selvamani** contributed the following: initial screening of different Raman lasers and characterization with Raman spectroscopy (support) and data analysis (support). **Joseph P. Smith** contributed the following: data analysis (support), conceptualization (support), and reviewing & editing the manuscript (support). **Richard Rustandi** contributed the following: in characterizing the viral particles effects before and after laser exposure (lead) and reviewing the manuscript (support). **Tiago Matos** contributed in the following: production and purification of viral material (lead), conceptualization, and reviewing manuscript (support). **Arezoo M. Ardekani** contributed the following: conceptualization (lead), writing – original draft (support), writing – formatting (support), writing – review & editing (support), project administration (lead), project coordination (support), and methodology (support). **Mohit S. Verma** contributed the following: conceptualization (lead), writing – original draft (support), writing – formatting (support), writing – review & editing (support), project administration (support), project coordination (lead), and methodology (lead).

## 8. Declaration of AI and AI-assisted technologies in the writing process

During the preparation of this work the authors used Grammarly (https://grammarly.com/) to check for grammar errors and improve the academic writing language. After using this tool/service, the authors reviewed and edited the content as needed and take full responsibility for the content of the publication.

