## Supporting Information for "Real-time monitoring of attenuated cytomegalovirus using Raman spectroscopy allows non-destructive characterization during flow"

Supplementary material:

**Section A:**

This section highlights the background controls and the internal standards used in this study. Figure S1 indicates the background Raman spectra for empty capillary and capillary filled with RO water. Table S1 enlists the calibration data collected before each experiment to ensure the optimum performance of the Raman Microscope.

Table S1:. Raman spectral analysis for Si reference as an internal standard reference

| Date | Curve name | Centre | Width | Height | % Gaussian | Type | Area | ChiSq |
| --- | --- | --- | --- | --- | --- | --- | --- | --- |
| 10/28/2022 | Curve 1 | 520.499 | 4.15 | 34073.7 | 27.98 | Mixed | 202150 | 0.31 |
| 11/8/2022 | Curve 1 | 520.484 | 3.98 | 34577.4 | 25.29 | Mixed | 198686 | 0.28 |
| 11/14/2022 | Curve 1 | 520.815 | 4.10 | 32963.9 | 26.95 | Mixed | 194260 | 0.30 |
| 11/16/2022 | Curve 1 | 520.664 | 4.05 | 33224 | 25.63 | Mixed | 193676 | 0.29 |
| 11/18/2022 | Curve 1 | 520.714 | 4.06 | 32791.3 | 25.15 | Mixed | 192026 | 0.29 |
| 11/22/2022 | Curve 1 | 520.267 | 4.08 | 32579.9 | 27.75 | Mixed | 190084 | 0.29 |
| 11/23/2022 | Curve 1 | 520.629 | 4.17 | 31389.1 | 29.04 | Mixed | 186216 | 0.32 |
| 11/30/2022 | Curve 1 | 520.531 | 4.11 | 33362.1 | 28.59 | Mixed | 195678 | 0.30 |
| 12/2/2022 | Curve 1 | 520.584 | 4.06 | 33305.7 | 26.04 | Mixed | 194738 | 0.29 |

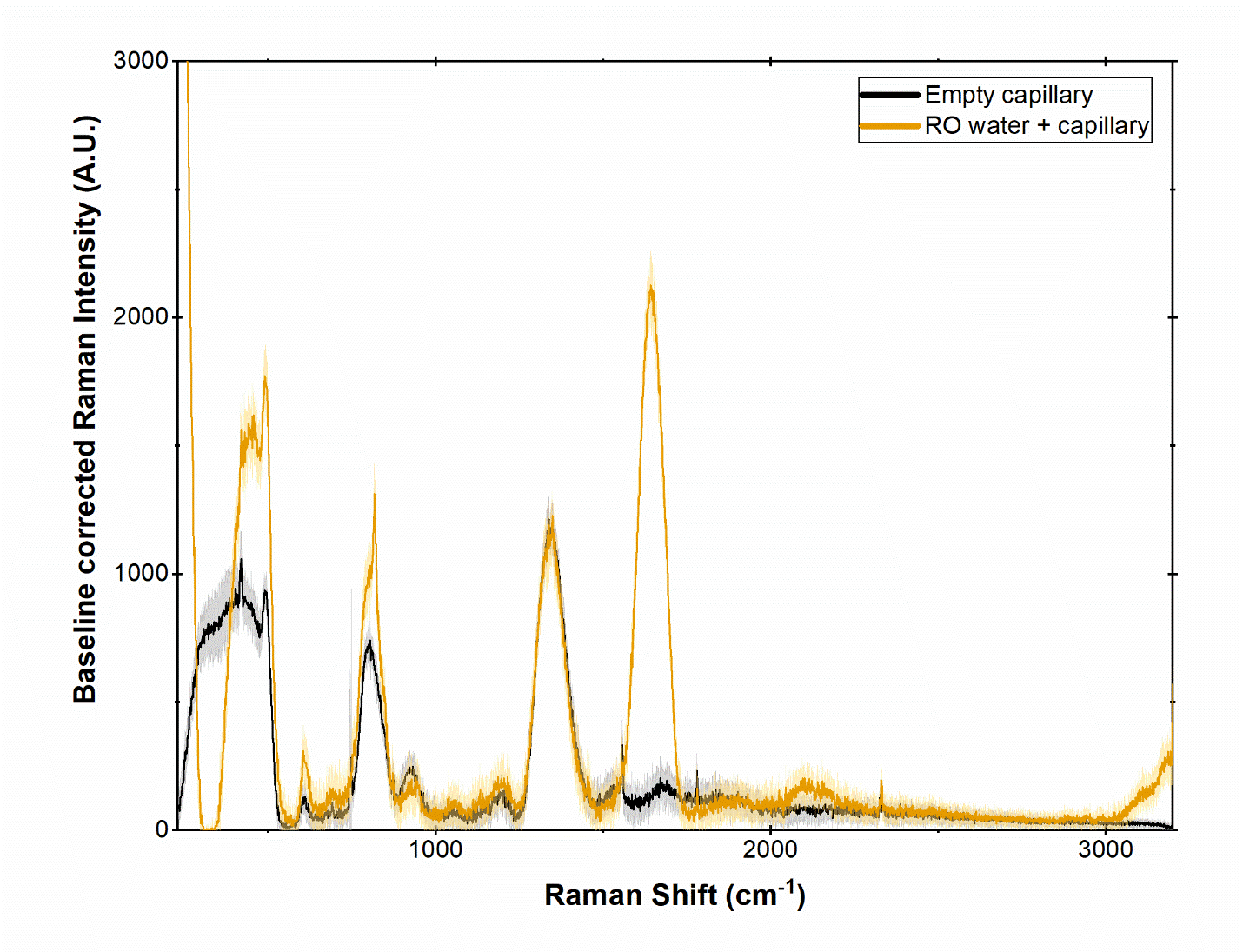

Figure S1: Baseline corrected Raman Spectra for Background (Empty capillary and RO water+ capillary).

**Section B:**

In this section, we provide the LoD calculations for other peaks from CMV viral particles that were distinct from the background peaks.

Table S2: LoD calculations for Raman peaks for CMV particles

| Peak (cm^-1^) | Linear Fit | R-square | LoB (Area Under the curve) | LoD (Area Under the Curve) | LoD (concentration, particles/mL) |
| --- | --- | --- | --- | --- | --- |
| 836 | y = 4.7E-11x + 16.58 | 0.76 | 14.12 | 20.74 | 8.84 x10^10^ |
| 921 | y = 1.59E-11x + 1.64 | 0.99 | 1.99 | 2.83 | 7.46 x10^10^ |
| 1072 | y = 7.8E-11x + 2.24 | 0.99 | 2.41 | 3.63 | 1.79 x10^10^ |
| 1127 | y = 6.06E-11x + 1.8 | 0.998 | 2.63 | 3.15 | 2.23 x10^10^ |
| 1463 | y = 2.16E-11x + 2.01 | 0.996 | 2.15 | 2.72 | 3.30 x10^10^ |
| 2900 | y = 5E-11x + 1.58 | 0.995 | 2.33 | 2.75 | 2.36 x10^10^ |

**Section C:**

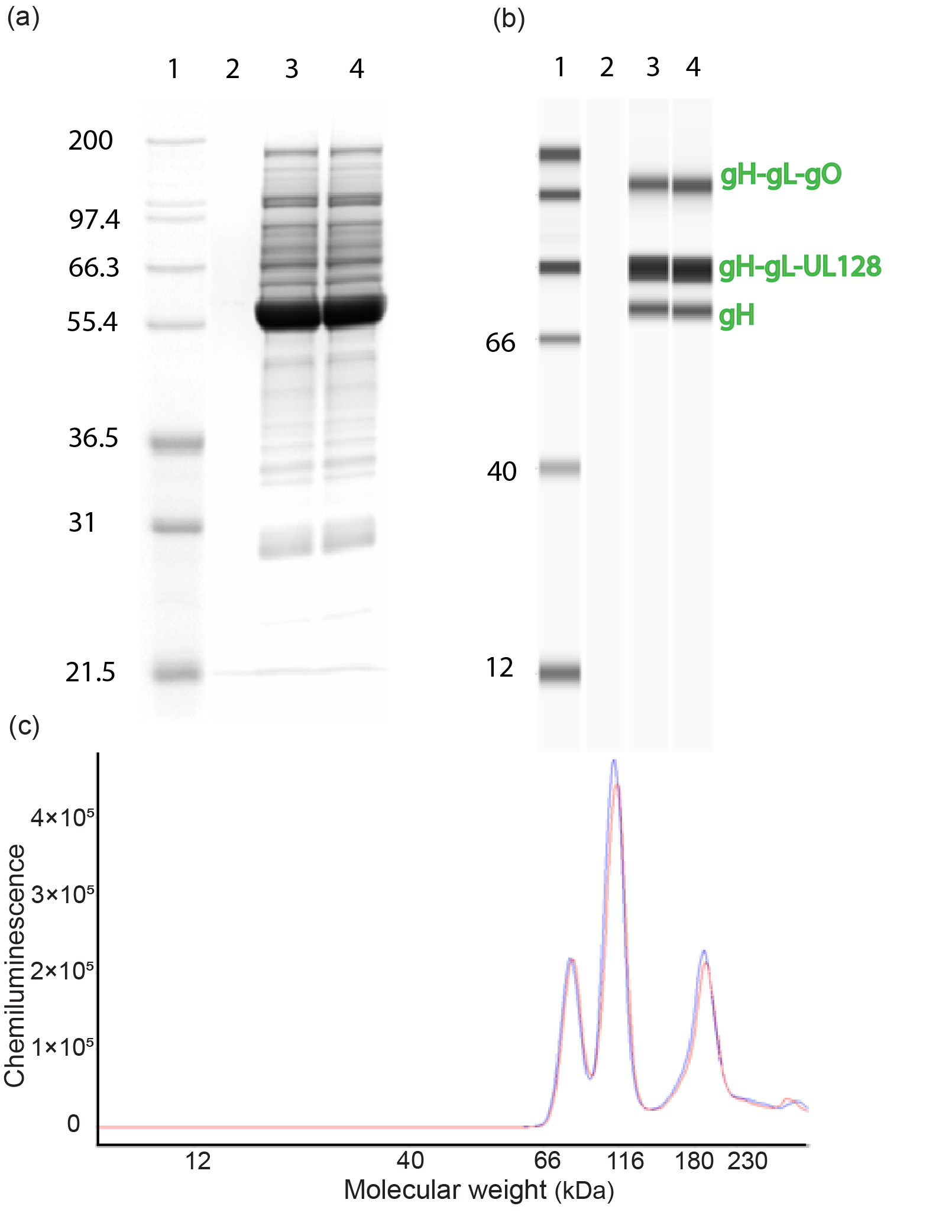
To study the functionality of the proteins before and after the laser exposure, a non-reducing Western blot was performed using the SimpleWestern system (ProteinSimple, San Jose, CA). The technology was previously described in detail.^1^ Briefly, the two samples were diluted 4.3x in a sample buffer containing SDS and iodoacetamide and heated at 70 °C for 10 min. Then, automated SDS PAGE gel and Western blot were run. The critical CMV antigens, pentameric (gH-gL-UL128) and trimeric (gH-gL-gO) proteins, were probed using specific anti-gH, and chemiluminescence signal was collected. ^2^

Figure S2: Functional analysis of the viral particles before and after laser exposure (wavelength: 785nm, Laser power: 0.5%(~1.5mW), power: 0.5%(~1.5mW), exposure time: 30s, n=1). (a) SDS page before and after laser exposure. Lane 3 – before laser exposure and Lane 4 - after laser exposure. (b) & (c) Western blot analysis and electrogram, respectively, for the samples b.

**Section D:**

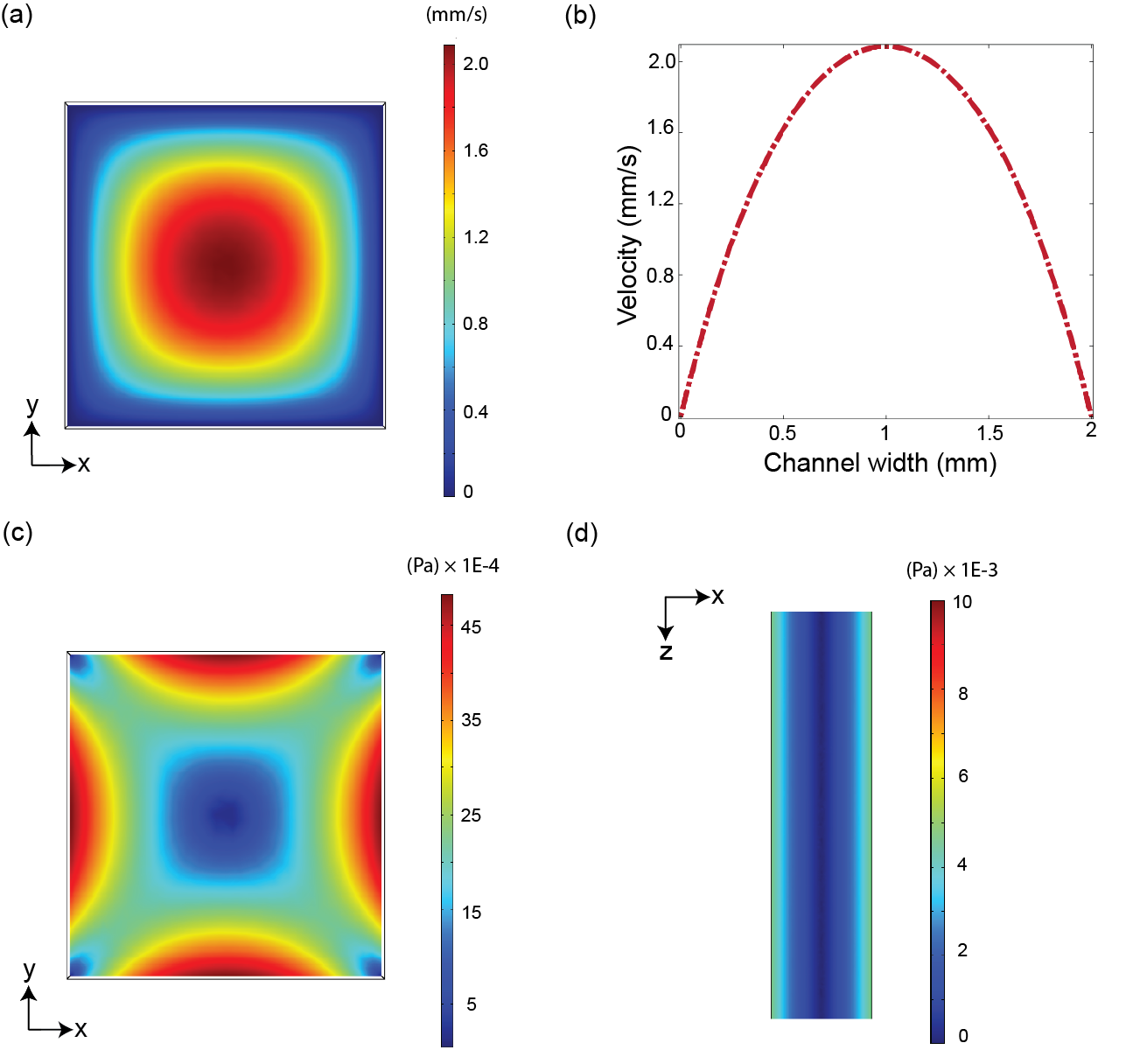
COMSOL Multiphysics 5.6 has been used to calculate the flow velocity and shear stress within the capillary tube. The flow inside the square capillary is considered steady-state 3D incompressible flow where the viscosity force is dominant, and Reynolds number is low$, Re=\frac{\rho UW}{\mu}\sim O($1$)$ ($\rho={10}^{3} (\frac{kg}{m^{3}})$ as density, $U=\left( 0.2-1 \right)\times{10}^{-3} (\frac{m}{s})$,$W=2\times{10}^{-3} (m)$, and $\mu={10}^{-3}(kg/ms$)). Also, we calculated the Stokes number (Stk=$\frac{\rho_{p}d_{p}^{2}U_{m}}{18W\mu}$), where $\rho_{d}\sim1700 (kg/m^{3})$ is viral particle density, $d_{p}\sim150 (nm)$ viral particle diameter, $U_{m}\sim\left( 0.4-2 \right)\times{10}^{-3} (\frac{m}{s})$ is maximum flow velocity,$W=2\times{10}^{-2} (m)$ is the channel width, and $\mu={10}^{-3}(kg/ms$) is the viscosity of the medium).

Figure S3: Velocity and shear stress calculation in the square capillary tube. (a) Velocity contour in the x-y plane. (b) Velocity profile (V_z) in the middle of the capillary in the z-direction. (c): Shear stress in the x-y plane. (d) Shear stress in the x-z plane.
